# Family history and APOE4 risk for Alzheimer’s Disease impact the neural correlates of episodic memory by early midlife

**DOI:** 10.1101/102699

**Authors:** M. N. Rajah, L. M. K. Wallace, E. Ankudowich, E. H. Yu, A. Swierkot, R. Patel, M. M. Chakravarty, D. Naumova, J. Pruessner, R. Joober, S. Gauthier, S. Pasvanis

**Author notes:** Corresponding author: M. Natasha Rajah, Room 2114 CIC Pavilion, Douglas Mental Health University Institute, 6875 LaSalle Blvd, Montreal, QC, Canada H4H 1R3.

## Abstract

Episodic memory impairment is a consistent, pronounced deficit in pre-clinical stages of late-onset Alzheimer’s disease (AD). Individuals with risk factors for AD exhibit altered brain function several decades prior to the onset of AD-related symptoms. In the current event-related fMRI study of spatial context memory we tested the hypothesis that middle-aged adults (MA; 40-58yrs) with a family history of late onset AD (MA_+FH_), or a combined +FH and apolipoprotein E ε4 allele risk factors for AD (MA_+FH+APOE4_), will exhibit differences in encoding and retrieval-related brain activity, compared to – FH –APOE4 MA controls. We also hypothesized that the two at-risk MA groups will exhibit distinct patterns of correlation between brain activity and memory performance, compared to controls. To test these hypotheses we conducted multivariate task, and behavior, partial least squares analysis of fMRI data obtained during successful context encoding and retrieval. Our results indicate that even though there were no significant group differences in context memory performance, there were significant differences in brain activity and brain-behavior correlations involving hippocampus, inferior parietal cortex, cingulate, and precuneus in MA with AD risk factors, compared to controls. In addition, we observed that brain activity and brain-behavior correlations in anterior-medial PFC and in ventral visual cortex differentiated the two MA risk groups from each other, and from MA_controls_. Our results indicate that functional differences in episodic memory-related regions are present by early midlife in adults with +FH and +APOE-4 risk factors for late onset AD, compared to middle-aged controls.

## 1. Introduction

Aging is associated with episodic memory decline: a reduced ability to encode, store and retrieve information about past events (recognition memory) in rich spatial and temporal contextual detail (context memory)[1–4]. These deficits negatively impact older adults’ quality of life [5] and can be an early sign of late-onset Alzheimer’s disease (AD)[6–9]. One promising way to support healthy brain aging and memory function into late life and prevent/delay AD onset is early identification of episodic memory decline in adults at risk of developing AD, and early intervention to prevent/delay further decline. To achieve these goals it is important to identify when episodic memory decline arises in adulthood, and determine how known risk factors for AD, e.g. having a family history of AD (+FH) or having an apolipoprotein E ε4 allele (+APOE4), alter memory and brain function at this critical time.

Recent studies of healthy adults show that episodic memory decline can be detected by early midlife (40 – 58 yrs) when memory is assessed using spatial context memory tasks [10–12]. In contrast, item recognition memory remains intact in early midlife[13]. This suggests that spatial context memory tasks are sensitive to detecting early episodic memory decline in healthy adults. Spatial context memory tasks require subjects to form item-location associations. As such, they are a type of associative memory task and place greater demands on recollection processes compared to item recognition tasks[14]. Neuroimaging studies of healthy young adults indicate that recollection of spatial contextual details relies on the activation of a distributed network of brain regions that include the medial temporal lobe (MTL), prefrontal cortex (PFC) and inferior parietal cortex [15–18]. Recently, we have examined the functional brain differences associated with context memory decline in middle-aged adults (MA) [12]. We reported group differences in ventrolateral prefrontal cortex (VLPFC) activity at encoding, and in ventral visual activity at retrieval, in MA compared to young adults. These activation differences were related to spatial context memory decline in MA and may reflect functional decline at midlife. In addition, MA also exhibited increased activity in anterior PFC at retrieval, compared to young adults, which correlated with better memory performance (potentially a mechanism for functional compensation).

Taken together, these findings indicate that associative spatial context memory tasks are a powerful tool for detecting behavioral and brain differences in episodic memory at early midlife. Thus, fMRI studies of spatial context memory can help us detect brain differences in *early* MA at risk of AD, compared to controls, which may be indicative of early pathological brain changes within the episodic memory system. Yet, the majority of fMRI studies of MA at risk of AD have used item recognition tasks. These studies have reported functional differences in brain regions in the MTL [19–22], inferior parietal cortex [23, 24], prefrontal cortex [25], and posterior midline cortex [21, 26]. However, the results are varied. For example, some of these studies report reduced brain activity in hippocampus and other areas, in at-risk groups vs. controls [19, 27, 28]; others report increased activity [20, 21]. Moreover, it is unclear whether these results were directly linked to episodic memory performance in MA with vs. without AD risk factors.

The goal of the current study was to understand the impact of having AD risk factors on the functional neural correlates of spatial context memory in early midlife (ages 40 – 58). Specifically, we conducted an event-related fMRI study in which the following MA groups were scanned while performing easy and hard versions of spatial context memory tasks: 1) -FH, -APOE4 MA (MA_controls_), 2) +FH, -APOE4 MA (MA_+FH_), and 3) +FH, +APOE4 MA (MA_+FH+APOE4_). Subjects were scanned during both encoding and retrieval phases of the memory tasks. The rationale for testing easy and hard versions of the task was to differentiate between performance effects, and group-by-performance interactions in brain activity. The rationale for scanning subjects during both encoding and retrieval was to examine group similarities and differences in phase (encoding/retrieval)-related activity and to understand how activity patterns at encoding related to those observed at retrieval, and vice versa. Examining brain activity during both encoding and retrieval, together, is important for determining whether group differences in regional activation are general across phases, and reflect general changes in regional brain function; or, if they are phase-specific and reflect group differences in task orientation and processes specific to encoding or retrieval. We hypothesized that having +FH or combined +FH, +APOE4 risk factors for AD would be related to differences in event-related activity in the MTL during encoding [21, 29]. We also hypothesize more general group differences in brain activity in other regions related to recollection-related processing, which are also implicated in the AD, i.e. inferior parietal cortex and PFC. To test these hypotheses we used multivariate “task” partial least squares analysis (T-PLS), a powerful method that allows one to identify whole-brain patterns of activity which maximally account for the co-variance between event-related brain activity and the experimental design [30]. We also hypothesized that having +FH and/or combined +FH, +APOE4 risk factors for AD would alter the correlation between brain activity in the aforementioned areas, and behavior. To test this hypothesis we used behavior-PLS (B-PLS). The current study is novel in that it is the first to use a spatial context memory tasks and multivariate PLS methods to assess functional brain differences in recollection-related brain activity at encoding *and* retrieval across at early middle-aged adults with +FH and with combined +FH, +APOE4, compared to controls.

## 2. Materials and Methods

### 2.1 Subjects

Fifty-one middle aged adults (MA; age range 41-58 yrs, mean age = 50.69 yrs, 40 females [10 menopausal, 4 on hormone replacement therapy]) were recruited using newspaper and online advertisements in Montreal, Canada. All subjects were healthy at the time of testing and had no history of neurological or psychiatric illness. All subjects were right-handed as measured by the Edinburgh Inventory for Handedness [31]. The study was approved by the Institutional Review Board of the Faculty of Medicine, McGill University, and all subjects provided informed consent to undergo neuropsychological testing, fMRI testing and to have their blood drawn for APOE genotyping.

#### 2.1.1 Neuropsychological assessment and exclusionary criteria

We administered the following battery of neuropsychological tests to screen out individuals suffering from psychiatric symptoms and cognitive impairment, and to obtain measures of memory and language function: Mini Mental Status Exam [MMSE, exclusion cut-off score < 27, [32]] the Beck Depression Inventory (BDI) [inclusion cut-of < 15 [33]], the American National Adult Reading Test (NART) [inclusion cut-off ≤ 2.5 SD for age and education [34]]. Additional medical exclusion criteria included having a history of mental health or substance abuse issues, neurological insult resulting a loss of consciousness > 5 min, diabetes, having untreated cataracts and glaucoma, smoking > 40 cigarettes a day; and having a current diagnosis of high cholesterol levels and/or high blood pressure left untreated in the past six months. All subjects who participated in the fMRI scanning session met these cut-off criteria. In addition, the California Verbal Learning Task (CVLT) was administered to assess item memory.

#### 2.1.2 Definitions of risk factors

Having a family history of late onset sporadic AD (+FH) was defined using the criteria used in the Cache County study [35, 36]: having a first degree relative, living or deceased, with a probable or confirmed diagnosis of AD. Having no family history of AD (–FH) was defined as the absence of first and second degree relatives with AD type dementia[36]. For APOE genotyping, genomic DNA was extracted from whole blood using the FlexiGene DNA kit from Qiagen (Qiagen, Ontario, Canada). Samples were genotyped with Sequenom iPLEX Gold Assay technology at Genome Quebec Innovation Centre (Quebec, Canada, [37]). APOE genotype results were used to stratify participants into three risk groups based on family history and genotype combination: no family history with APOE ε3/3 genotype (MA_controls_), family history with APOE ε3/3 (MA_+FH_), and family history with APOE ε3/4 (MA_+FH+APOE4_ 4).

### 2.2 Experimental Protocol

#### 2.2.1 Cognitive activation task

Subjects performed easy and difficult versions of spatial context memory tasks while undergoing blood-oxygen-level-dependent (BOLD) fMRI scanning. The rationale for including easy and difficult versions of the task was to allow for examination of performance effects and group^*^ performance interactions in behavior and brain activity. Subjects were scanned during encoding and retrieval. The rationale for scanning during both encoding and retrieval was so that we could directly examine how activation related successful encoding related to activity during successful retrieval. This allowed us to identify areas that were similarly activated during the two memory phases across groups, and to explore group differences in memory phase-related modulation. E-Prime (Psychology Software Tools, Inc.; Pittsburgh, PA, USA) was used to present memory tasks and collect behavioural data (accuracy and reaction time)

##### Spatial context encoding

Subjects were shown black and white photographs of human faces, presented one at a time on either the left or right side of a monitor, for 2 sec each, with a variable inter-trial interval (ITI) of 2.2 – 8.8 sec (mean ITI 5.13 sec). Subjects were instructed to rate whether the face was pleasant/neutral using a button press, and to encode the spatial location (left/right) in which the face was presented. During easy spatial context memory tasks (SE) subjects encoded six face stimuli, and during hard spatial context memory asks (SH) subjects encoded 12 face stimuli. Subjects were aware at encoding that their memory for spatial location would be tested following a 1 min break. During the break subjects performed a verbal alphabetizing distractor task to prevent rehearsal of face stimuli, and to ensure retrieval involved long-term, episodic memory processes. There were 12 blocks of SE encoding blocks and 6 blocks of SH encoding tasks. There were 72 encoding stimuli per task.

##### Spatial context retrieval

Subjects were presented with two previously encoded face stimuli for 6 sec, with variable ITI (as stated above), and were asked to select which face was originally presented on the left (or the right) side of the screen at encoding using a button press, depending on the retrieval cue. Thus, subjects had to recollect the spatial location of the encoded face to perform the task above chance. There were 12 SE retrieval blocks and 6 SH retrieval blocks. There were 36 retrieval stimuli per task.

#### 2.2.2. Behavioral Data Analysis

SPSS for Windows (version 17.0) was used to conduct between group one-way ANOVAs on demographic and neuropsychological variables to ensure groups were matched on age, education, and neuropsychological tests. In addition, 3 (group: MA_controls_; MA_+FH_; MA_+FH+APOE4_) × 2 (event-type: SE, SH) repeated measures ANOVAs were conducted to examine group main effects, task main effects and group^*^task interactions in spatial context memory accuracy (% correct) and reaction time (RT; msec) during easy and hard task versions (significance threshold p < 0.05). Post-hoc Tukey’s tests were conducted on the group variable to clarify any significant group main effects and interaction effects.

### 2.2.2. MRI Data Acquisition

Structural and functional magnetic resonance images were acquired using a 3T Siemens Trio scanner, located at the Douglas Brain Imaging Centre. Subjects lay supine in the scanner wearing a standard head coil. T1-weighted structural images were acquired at the beginning of the fMRI session using a 3D gradient echo MPRAGE sequence (acquisition time: 5 min, 3sec; TR=2300 msec TE=2.98 msec, flip angle=9 degrees, 176 1mm saggital slices, 1 × 1 × 1 mm voxels, FOV=256mm2). BOLD images were acquired using a single-shot T2^*^-weighted gradient echo-planar imaging (EPI) pulse sequence (TR=2000ms, TE=30ms, FOV=256mm2, matrix size=64 × 64, in-plane resolution 4 × 4 mm, 32 oblique 4mm slices with no slice gap) while participants performed the memory tasks. A mixed rapid event-related design was used with variable ITI (as stated above) to add jitter to the event-related acquisitions.

Visual task stimuli were presented on a computer using E-Prime (described above) and were back-projected onto a screen in the scanner bore. The screen was visible to participants lying in the scanner via a mirror mounted within the standard head coil. Participants requiring correction for visual acuity wore plastic corrective glasses. A fiber-optic 4-button response box was used by subjects to make task-related responses.

### 2.4. MRI Data Analysis

#### 2.4.1. Functional MRI Analysis

##### Preprocessing

Images were reconstructed from raw (k-space), converted to ANALYZE format, and preprocessed using Statistical Parametric Mapping (SPM) version 8 software (http://www.fil.ion.ucl.ac.uk/spm) run with MATLAB (www.mathworks.com) on a Linux platform. Images acquired during the first 10 sec of scanning were removed from analysis to ensure all tissue had reached steady state magnetization. All functional images were realigned to the first image and corrected for movement artifacts using a 6 parameter rigid body spatial transform. If a subject had more than 4mm movement, they were discarded from analysis. None of the subjects included in the current study moved more than 4 mm and the total sample was included in all analyses.

Functional images were then spatially normalized using the MNI EPI-template (available in SPM) at 4mm^3^ voxel resolution using and default settings in SPM8 12 parameter affine transformation, as follows: template weighting = 0, affine regularisation to the ICBM/MNI space template, nonlinear frequency cutoff =25, default nonlinear iterations =16, nonlinear regularisation = 1. Writing options were set to preserve concentrations, 2X 3 double bounding box, 4×4×4 voxel size, and trilinear interpolation. Images were smoothed using an 8mm full-width half-maximum (FWHM) isotropic Gaussian kernel. ArtRepair toolbox for SPM8 was used to correct for bad slices prior to realignment and for bad volumes after normalization and smoothing (http://cibsr.stanford.edu/tools/human-brain-project/artrepair-software.html).

##### Multivariate fMRI Data Analysis

Spatio-temporal Partial Least Squares (PLS) was used to conducted event-related fMRI data analysis using PLSGUI software (https://www.rotman-baycrest.on.ca/index.php?section=84). This multivariate approach was chosen due to its ability to detect spatially and temporally distributed patterns of activated voxels that differ across experimental conditions and/or relate to a specific behavioral measure. (McIntosh, Chau, & Protzner, 2004). Details on this method have been published elsewhere (Krishnan, Williams, McIntosh, & Abdi, 2011; McIntosh & Lobaugh, 2004). Mean centered Task PLS (T-PLS) was used to examine group similarities and differences in patterns of whole brain activity related to encoding and retrieval during SE and SH tasks [30]. Behavior PLS (B-PLS) was used to examine group similarities and differences in patterns of whole brain activity *that were directly correlated to* SE and SH retrieval accuracy. For both T-PLS and B-PLS analyses, fMRI analysis was restricted to correctly encoded and correctly retrieved (successful) events.

For both T-PLS and B-PLS analyses, the first step was to represent the fMRI data for correctly encoded and retrieved events in an fMRI data matrix. PLS for event-related fMRI converts the three dimensional event-related fMRI data to a two-dimensional data matrix by ‘flattening’ the temporal dimension (t), so that time series of each voxel (m) is stacked side-by-side across the columns of the data matrix (column dimension = m^*^t). The rows of the 2D data matrix for an individual subject represents activity in each event stacked within experimental condition (row dimension = events^*^conditions). The event-related data for each event-type was averaged within subject, and subjects’ fMRI data were stacked within group, and groups were stacked above one another [30]. The stacked data matrix contained the fMRI data for each event onset (time lag =0) as well as the subsequent seven TRs/time lags (TR = 2 sec^*^ 7 = 14 sec) following event onset for *successfully* encoded (eSE, eSH) and *successfully* retrieved (rSE and rSH) events were stacked by subject within group to create a between group fMRI data matrix. All subjects analyzed had a minimum of 17 correct events per event type (mean *#* of correct events for SE task = 30 within each group; and for SH task = 29 for MA_controls_ MA_+FH_ and 30 for MA_+FH+APOE4_).

In T-PLS, this fMRI data matrix was mean centered, column-wise, within each event type, and underwent singular value decomposition (svd). SVD yields a set of mutually orthogonal latent variables (LVs) in descending order of the magnitude of covariance explained. The number of LVs produced is equivalent to the number of event/task types included in the analysis^*^ the number of groups; in this analysis there were 12 (4 event-types^*^ 3 groups). Each mean centered T-PLS LV consists of: i) a singular value, reflecting the amount of covariance accounted for by the LV; ii) design saliences, representing the event-related contrast effect identified by the LV that are presented as design salience plot in the results, and are similar to contrast effects identified in univariate fMRI analysis; and, iii) a singular image (s.i.) that represents a pattern of event-related brain activity which corresponds to the design salience plot. Importantly, the pattern of whole brain activity identified in the singular image symmetrically relates to the design salience plot. The singular image includes brain saliences, which are numerical weights assigned to each voxel at each TR/time lag included in the data matrix. Brain saliences can be negative or positive. Brain regions with positive voxel saliences were positively related to the experimental effect identified by the design salience plot, and those with negative voxel saliences were negatively related to this effect.

In B-PLS, the between group fMRI data matrix was correlated with a behavioral vector containing the mean retrieval accuracy for SE and SH tasks (% correct spatial context retrieval), stacked in the same order as the data matrix (subject within group). SVD of this cross-correlation matrix was conducted to yield a series of LVs. The output is similar to the T-PLS output, but instead of design saliences, the B-PLS analysis yields: i) a singular value, ii) a singular image consisting of positive and negative brain saliences, and iii) *a correlation profile* depicting how subjects’ retrieval accuracy correlates with the pattern of whole brain activity identified in the singular image. The correlation profile and voxel saliences represent a symmetrical pairing of i) brain-behavior correlation patterns for each group to ii) a pattern of whole brain activity, respectively. As with the T-PLS analysis, brain saliences can have positive or negative values, and reflect whether activity in a given voxel is positively or negatively associated with the correlation profile depicted.

Significance testing of LVs identified from the T-PLS and B-PLS was conducted using 1000 permutation tests, respectively. For T-PLS the permutation test involved sampling without replacement to reassign the event/condition order within subject. For each permuted iteration a PLS was recalculated, and the probability that the permuted singular values exceeded the observed singular value for a given LV was used to assess significance at p<0.05[30]. The permutation method used met the exchangeability criterion as described in [38]. The permutation method for B-PLS was identical except that the permutation test involved reassignment of the link between subjects’ behavioral measure (retrieval accuracy) - event/condition within subject.

In addition, for each analysis, the standard error of brain saliences for each LV was determined by conducting 500 bootstraps using sampling of subjects with replacement, while maintaining the experimental event/condition order for all subjects [39]. The ratio of the original brain salience to bootstrap standard error (bootstrap ratio; BSR), which is similar to a z-score [38], was calculated for each voxel-based brain salience. The bootstrap result identifies the maximal reliable patterns of positive and negative brain saliences represented by the singular image. A threshold of BSR ≥ ±3.5, p < 0.001, with a minimum spatial extent = 10 voxels, was used to report the stable maximal brain saliences identified in each significant LV. The minimal spatial extent = 10, was selected so that our reported results were comparable to prior work using both univariate and multivariate methods [40]. The BSR of a significant voxel salience reflects the stability of its activation.

We also computed temporal brain scores for each task in each significant LV. Temporal brain scores represent the degree to which each subject expresses the pattern of brain activity identified by the s.i., in relation to its paired design salience (T-PLS)/correlation profile (B-PLS), at each time lag. The temporal brain score was used to identify the time lags at which the LV effect was maximally differentiated within the temporal window sampled[38]. We only report reliable (BSR ≥ ±3.5) brain saliences from those time lags [41, 42]. In the current analyses the peak time lags were lags 2 – 5 (4 – 10 sec post event-onset). The peak coordinates for reliable brain saliences were converted to Talairach space using the icbm2tal transform (Lancaster et al. 2007) as implemented in GingerAle 2.3 (Eickhoff et al. 2009). Since our acquisition incompletely acquired the cerebellum, peak coordinates from this region were not reported. The Talairach and Tournoux atlas (1988) was used to identify the Brodmann area (BA) localizations of significant activations.

## 3. Results

### 3.1. Demographics and Behavioral Results

Of the 51 MA tested, 26 were identified as MA_controls_ (-FH, -APOE4); 14 MA were MA_+FH_ (- APOE4); and 11 were MA_+FH+APOE4_. Table 1 presents the demographic, neuropsychological and behavioral data from the fMRI tasks for each group. There were no significant group differences in any of the demographic or neuropsychological variables. The group (3) x task (2) repeated measures ANOVA for retrieval accuracy (% correct) revealed no significant effects, although there was a trend towards there being a task main effect (p = 0.10) due to accuracy on SH tasks being lower than on SE tasks. The group (3) x task (2) repeated measures ANOVA for reaction time (RT, msec) revealed a significant task main effect (F_1,48_=12.85, p < 0.001, SH RT slower than SE), but no other main effects or interactions. Therefore, the three groups were matched on task performance, and retrieval performance on SH tasks was worse than retrieval performance on SE tasks.

**Table 1.** Demographics and Behavioral Data

|  | Middle-aged adults with no family history of dementia (-FH), with APOE e3/3 genotype | Middle-aged adults with a family history of Alzheimer's disease (+FH), with APOE e3/3 genotype | Middle-aged adults with a family history of Alzheimer's disease (+FH), with APOE e3/4 genotype |
| --- | --- | --- | --- |
| Sample size (n) | 26 | 14 | 11 |
| Age (mean ± SE) | 49.27 ± 1.13 | 51.43 ± 1.08 | 51.91 ± 1.52 |
| Gender (n, [%] female) | 21 [81] | 10 [71] | 9 [82] |
| Education (years, mean ± SE) | 15.46 ± 0.37 | 15.64 ± 0.43 | 14.92 ± 0.62 |
| BDI (mean ± SE) | 3.85 ± 0.85 | 4.64 ± 1.33 | 5.41 ± 1.33 |
| CVLT: delayed free recall (mean ± SE) | 12.35 ± 0.46 | 12.86 ± 0.61 | 12.64 ± 0.66 |
| CVLT: delayed cued recall (mean ± SE) | 13.04 ± 0.41 | 13.14 ± 0.51 | 12.73 ± 0.82 |
| CVLT: delayed recognition (mean ± SE) | 15.19 ± 0.15 | 15.07 ± 0.27 | 14.73 ± 0.56 |
| SE reaction time (msec; mean ± SE) | 2467.40 ± 77.36 | 2485.71 ± 135.01 | 2566.09 ± 200.67 |
| SH reaction time (msec; mean ± SE) | 2578.68 ± 61.55 | 2649.64 ± 135.36 | 2672.19 ± 131.08 |
| SE retrieval accuracy (% correct; mean ± SE) | 0.86 ± 0.02 | 0.86 ± 0.03 | 0.85 ± 0.03 |
| SH retrieval accuracy (% correct; mean ± SE) | 0.81 ± 0.03 | 0.87 ± 0.02 | 0.83 ± 0.03 |
| Left HC Volume (mm; mean ± SE) | 2807 ± 75 (N = 25) | 2875 ± 440 (N = 13) | 2750 ± 113 (N = 11) |
| Right HC Volume (mm; mean ± SE) | 2906 ± 80 (N = 25) | 2972 ± 396 (N = 13) | 2885 ± 114 (N = 11) |

Note: This table presents the group means and standard errors (S.E.) for demographic, neuropsychological, fMRI behavioral measures and hippocampal (HC) volumes obtained. BDI = Beck Depression Inventory; CVLT = California Verbal Learning Task. All CVLT results presented are “hits”. There were no significant group differences on these measures.

**Table 2:** Task PLS Results - LV 1

| Lag | BSR | Spatial Extent | Talairach Coordinates (mm) |  |  | Gyrus Location | Brodmann Area (BA) |
| --- | --- | --- | --- | --- | --- | --- | --- |
|  |  |  | X | Y | Z |  |  |
| Positive Saliences: Regions that were more active during retrieval vs. encoding across all groups |  |  |  |  |  |  |  |
| Left Hemisphere |  |  |  |  |  |  |  |
| 2,3,4 | 9.13 | 76 | -31 | 21 | 2 | Inferior Frontal Gyrus/Insula | BA 47 |
| 2,3 | 5.41 | 107 | -38 | -78 | 14 | Middle Occipital Gyrus | BA 19 |
| 2 | 5.23 | 27 | -16 | 2 | 8 | Putamen |  |
| 2 | 5.09 | 88 | -13 | -59 | 16 | Posterior Cingulate | BA 23 |
| 3 | 6.32 | 806 | -5 | -10 | 54 | Precentral Gyrus | BA 6 |
| 3 | 5.31 | 44 | -38 | 11 | 30 | Middle Frontal Gyrus | BA 9/44 |
| 3 | 4.49 | 14 | -12 | -1 | 0 | Lentiform Nucleus |  |
| 3,4 | 5.19 | 69 | -31 | -80 | 32 | Superior Occipital Gyrus | BA 19 |
| 4 | 5.26 | 60 | -46 | -73 | 0 | Inferior Temporal Gyrus | BA 37 |
| 4 | 5.23 | 32 | -38 | -52 | -16 | Fusiform Gyrus | BA 37 |
| 4 | 4.11 | 32 | -42 | -35 | 40 | Inferior Parietal Lobule | BA 40 |
| 4 | 4.08 | 11 | -38 | -6 | 10 | Insula |  |
| 4 | 4.05 | 16 | -53 | -26 | 19 | Postcentral Gyrus | BA 43 |
| Right Hemisphere |  |  |  |  |  |  |  |
| 2,3 | 10.24 | 114 | 29 | 25 | 0 | Insula |  |
| 2 | 5.26 | 28 | 14 | 5 | 9 | Putamen |  |
| 2 | 4.39 | 57 | 28 | -75 | 19 | Middle Occipital Gyrus | BA 19 |
| 3,4 | 7.48 | 423 | 10 | -56 | 21 | Posterior Cingulate | BA 30 |
| 3 | 5.41 | 65 | 39 | 10 | 31 | Middle Frontal Gyrus | BA 9/44 |
| 4 | 6.06 | 143 | 32 | 24 | 4 | Inferior Frontal Gyrus | BA 45 |
| 5 | 4.75 | 59 | 36 | -45 | -17 | Fusiform Gyrus | BA 37 |
| Negative Saliences: Regions that were more active during encoding vs. retrieval across all groups |  |  |  |  |  |  |  |
| Left Hemisphere |  |  |  |  |  |  |  |
| 2 | -10.77 | 925 | -27 | -99 | -2 | Cuneus | BA 18 |
| 2 | -7.99 | 426 | -57 | -41 | 21 | Superior Temporal Gyrus | BA 22 |
| 2 | -6.09 | 137 | -42 | 22 | -5 | Inferior Frontal Gyrus | BA 47 |
| 2 | -5.72 | 124 | -12 | 48 | 34 | Superior Frontal Gyrus | BA 9 |
| 2 | -5.49 | 44 | -53 | -13 | 45 | Postcentral Gyrus | BA 3 |
| 2 | -4.24 | 27 | -35 | -18 | 16 | Insula |  |
| 3,4 | -8.02 | 590 | -60 | -54 | -2 | Middle Temporal Gyrus | BA 37 |
| 3 | -6.97 | 375 | -8 | 36 | 4 | Anterior Cingulate | BA 24 |
| 3,4,5 | -8.22 | 263 | -46 | -6 | 50 | Precentral Gyrus | BA 6 |
| 4 | -4.59 | 17 | -39 | 17 | 48 | Superior Frontal Gyrus | BA 8 |
| 5 | -5.15 | 24 | -5 | 1 | 51 | Medial Frontal Gyrus | BA 6 |
| 5 | -4.81 | 36 | -31 | 43 | 11 | Superior Frontal Gyrus | BA 10 |
| Right Hemisphere |  |  |  |  |  |  |  |
| 2 | -8.45 | 590 | 58 | -23 | 25 | Postcentral Gyrus | BA 2 |
| 2 | -6.52 | 92 | 51 | 8 | -23 | Superior Temporal Gyrus | BA 38 |
| 2 | -5.44 | 88 | 40 | 45 | 24 | Middle Frontal Gyrus | BA 10 |
| 2 | -4.06 | 12 | 36 | -9 | -3 | Clastrum |  |
| 3,4,5 | -14.12 | 1177 | 3 | -87 | -8 | Lingual Gyrus | BA 17,18 |
| 3,4 | -7.80 | 472 | 55 | -23 | -8 | Middle Temporal Gyrus | BA 21 |
| 3,4 | -6.23 | 192 | 32 | 20 | 50 | Superior Frontal Gyrus | BA 8 |
| 3,4,5 | -5.75 | 18 | 54 | -10 | 44 | Precentral Gyrus | BA 4 |
| 3 | -4.41 | 13 | 40 | 9 | -34 | Middle Temporal Gyrus | BA 21,38 |
| 4 | -6.24 | 12 | 50 | -48 | 47 | Inferior Parietal Lobule | BA 40 |
| 4 | -5.28 | 179 | 10 | 32 | 4 | Anterior Cingulate Gyrus | BA 24 |
| 5 | -5.38 | 21 | 25 | 47 | 6 | Superior Frontal Gyrus | BA 10 |

Note: Temporal lag represents the time after event onset, when a cluster of voxels exhibited a contrast effect of interest. The bootstrap ratio threshold was set to $\pm > 3.5$ , and identified dominant and stable activation clusters. The spatial extent refers to the total number of voxels included in the cluster (threshold = 10). Gyrus location and Brodmann areas (BAs) were determined by referring to Talairach and Tournoux (1988).

### 3.2. FMRI Results

We conducted two multivariate analyses to identify: 1) group similarities and differences in event-related *activity* during successful spatial context encoding and retrieval (T-PLS), and 2) group similarities and differences in the correlations between event-related activity and retrieval accuracy (B-PLS). These two analyses are complementary and results from both analyses need to be considered together in order to understand how having specific risk factors for AD impact brain activity in a behaviorally relevant manner. In the following sections we describe the T-PLS and B-PLS results separately. In the Discussion we focus on regions consistently identified across PLS methods to help clarify their roles in context memory encoding and retrieval in MA with vs. without AD risk factors.

#### 3.2.1. Task PLS Results

##### Group similarities in event-related brain activity

The mean centered T-PLS analysis identified three significant LVs. The first two LVs identified group similarities in task-related brain activity. Figures 1A and 1B present the design salience plot and singular image for these LVs. T-PLS LV1 (accounted for 45.95% cross-block covariance) and identified brain regions that were differentially activated during successful context encoding vs. retrieval in all three groups. Positive salience brain regions were more active during retrieval, compared to encoding, across all groups. Negative salience brain regions were more active during encoding, compared to retrieval, across all groups.

T-PLS LV2 (accounted for 14.44% cross-block covariance) identified brain regions that were differentially activated during SH encoding, compared to SE encoding in all three groups. Positive salience regions were more active during SH encoding, compared to SE encoding. Negative salience brain regions were more active during SE encoding, compared to SH encoding. The local maxima from these two LVs are presented in Tables 2 and 3. In general, this LV identified performance/difficulty-related effects at encoding which was common across groups.

##### Group differences in event-related brain activity

T-PLS LV3 (accounted for 12.81% cross-block covariance) identified a complex three-way interaction between group^*^task^*^phase (encoding/retrieval). This LV was of primary interest in this study, since it identified group differences in event-related brain activity. Figures 2A and 2B present the singular image and design salience plot for this LV. Table 4 presents the local maxima identified by T-PLS LV3. Positive brain salience regions from Table 4 were generally more active during encoding, compared to retrieval, in MA_controls_ and MA_+FH_. These regions included left angular gyrus, precuneus, and cingulate gyrus. Interestingly, these same regions were more active during SE retrieval, compared to SE encoding, in MA_+FH+APOE4_. Negative brain salience regions reflected the opposite effect. These regions were more active during retrieval, compared to encoding, in MA_controls_ and MA_+FH_. In contrast these same regions were more active during SE encoding, compared to SE retrieval in MA_+FH+APOE4_. Negative salience brain regions included bilateral fusiform cortices.

**Table 4:** Task PLS Results – LV 3

| Lag | BSR | Spatial Extent | Talairach Coordinates (mm) |  |  | Gyrat Location | Brodmann Area (BA) |
| --- | --- | --- | --- | --- | --- | --- | --- |
|  |  |  | X | Y | Z |  |  |
| *Positive Saliences: Regions that were more active during encoding vs. retrieval in MA control and MA+FH groups, but more active during retrieval vs. encoding of SE tasks in MA+FH+APOE4 group* |  |  |  |  |  |  |  |
| Left Hemisphere |  |  |  |  |  |  |  |
| 2,3, 4 | 6.33 | 174 | -54 | -58 | 38 | Angular Gyrus | BA 40 |
| 2 | 5.26 | 42 | -49 | -32 | 11 | Superior Temporal Gyrus | BA 22 |
| 2 | 4.27 | 21 | -9 | -23 | 63 | Precentral Gyrus | BA 6 |
| 4, 5 | 4.90 | 207 | -2 | -36 | 48 | Precuneus | BA 7 |
| 4 | 4.48 | 10 | -61 | -52 | 20 | Superior Temporal Gyrus | BA 22 |
| 4,5 | 4.75 | 36 | -16 | 10 | 37 | Cingulate Gyrus | BA 24 |
| 5 | 4.66 | 39 | -17 | -41 | 58 | Paracentral Lobule | BA 5 |
| Right Hemisphere |  |  |  |  |  |  |  |
| 2,3 | 6.13 | 56 | 54 | -47 | 44 | Inferior Parietal Lobule | BA 40 |
| 2 | 4.71 | 20 | 44 | 29 | -3 | Inferior Frontal Gyrus | BA 47 |
| 2 | 4.44 | 27 | 20 | -25 | 53 | Precentral Gyrus | BA 4 |
| 2,3,5 | 4.64 | 16 | 58 | -32 | 2 | Middle Temporal Gyrus | BA 21 |
| 4 | 4.24 | 32 | 21 | -17 | 43 | Cingulate Gyrus | BA 24 |
| 5 | 5.31 | 112 | 6 | -15 | 61 | Medial Frontal Gyrus | BA 6 |
| 5 | 4.90 | 19 | 18 | -2 | 1 | Putamen |  |
| 5 | 4.67 | 35 | 20 | -42 | 66 | Postcentral Gyrus | BA 5 |
| *Negative Saliences: Regions that were more active during retrieval vs. encoding in MA controls and MA+FH groups, but more active during encoding vs. retrieval of SE tasks in MA+FH+APOE4 group* |  |  |  |  |  |  |  |
| Left Hemisphere |  |  |  |  |  |  |  |
| 3 | -4.39 | 18 | -42 | -75 | -14 | Fusiform Gyrus | BA 19 |
| 4 | -4.53 | 22 | -27 | -95 | -5 | Lingual Gyrus | BA 18 |
| Right Hemisphere |  |  |  |  |  |  |  |
| 4 | -4.77 | 56 | 36 | -75 | -13 | Fusiform Gyrus | BA 19 |

Note: Temporal lag represents the time after event onset, when a cluster of voxels exhibited a contrast effect of interest. The bootstrap ratio threshold was set to $\pm > 3.5$ , and identified dominant and stable activation clusters. The spatial extent refers to the total number of voxels included in the cluster
(threshold = 10). Gyrus location and Brodmann areas (BAs) were determined by referring to Talairach and Tournoux (1988).

#### 3 2 2. B-PLS Results

##### Brain regions exhibiting group similarities in brain-behavior correlations at encoding, and group differences in brain-behavior correlations at retrieval

The B-PLS analysis identified two significant LVs. Figure 2A presents the singular image and the corresponding bar graph depicting the brain activity-behavior correlation profile for B-PLS LV1, which accounted for 32.36% of the cross-block covariance. Table 5 lists the local maxima from this LV.

**Table 5:** Behavior PLS Results – LV 1

| Lag | BSR | Spatial Extent | Talairach Coordinates (mm) |  |  | Gyrat Location | Brodmann Area (BA) |
| --- | --- | --- | --- | --- | --- | --- | --- |
|  |  |  | X | Y | Z |  |  |
| *Positive Saliences: Regions where there was a positive correlation between encoding activity and subsequent memory for all groups, and between retrieval activity during SE and accuracy for MA +FH; but, where there was a negative correlation between retrieval activity and retrieval accuracy in MA +FH +APOE4* |  |  |  |  |  |  |  |
| Left Hemisphere |  |  |  |  |  |  |  |
| 2,3 | 5.17 | 54 | -13 | -17 | 49 | Cingulate Gyrus | BA 24 |
| 3 | 5.48 | 122 | -24 | -11 | 64 | Precentral Gyrus | BA 6 |
| 3 | 5.40 | 14 | -45 | 16 | 12 | Inferior Frontal Gyrus | BA 44 |
| 3 | 4.54 | 24 | -50 | -19 | 27 | Postcentral Gyrus | BA 2 |
| 3 | 3.85 | 11 | -35 | -48 | 14 | Superior Temporal Gyrus | BA 22 |
| 4 | 5.54 | 128 | -13 | -29 | 55 | Paracentral Lobule | BA 40 |
| 4 | 5.43 | 17 | -34 | 33 | 0 | Inferior Frontal Gyrus | BA 45 |
| 4 | 5.33 | 44 | -42 | -47 | 46 | Inferior Parietal Lobule | BA 40 |
| 4,5 | 5.62 | 29 | -57 | -56 | 27 | Superior Temporal Gyrus | BA 39 |
| 4,5 | 6.31 | 1125 | -13 | -44 | 47 | Precuneus | BA 7 |
| 5 | 5.90 | 41 | -34 | -6 | -26 | Hippocampal Gyrus* | BA 35 |
| Right Hemisphere |  |  |  |  |  |  |  |
| 2 | 4.89 | 30 | 2 | -32 | 41 | Cingulate Gyrus | BA 31 |
| 3 | 4.22 | 10 | 62 | -52 | 18 | Superior Temporal Gyrus | BA 22 |
| 3 | 3.96 | 22 | 13 | -47 | 47 | Precuneus | BA 7 |
| 4 | 5.75 | 28 | 28 | -18 | 53 | Precentral Gyrus | BA 4 |
| 4 | 4.56 | 27 | 21 | 20 | 7 | Insula |  |
| 4,5 | 5.12 | 105 | 36 | -30 | 24 | Postcentral Gyrus | BA 40 |
| 4,5 | 4.32 | 15 | 58 | -50 | 33 | Supramarginal Gyrus | BA 40 |
| 5 | 7.96 | 93 | 9 | -10 | 54 | Cingulate | BA 24 |
| 5 | 4.70 | 35 | 43 | 8 | 17 | Inferior Frontal Gyrus | BA 44 |
| 5 | 4.21 | 11 | 40 | 45 | 24 | Middle Frontal Gyrus | BA 10 |
| 5 | 4.02 | 10 | 32 | 25 | 40 | Middle Frontal Gyrus | BA 8 |
| *Negative Saliences: Regions where there was a negative correlation between encoding activity and subsequent memory for all groups, and between retrieval activity during SE and accuracy for MA +FH; but, where there was a positive correlation between retrieval activity and retrieval accuracy in MA +FH +APOE4* |  |  |  |  |  |  |  |
| Left Hemisphere |  |  |  |  |  |  |  |
| 3 | -5.47 | 105 | -8 | 14 | -5 | Caudate |  |
| 5 | -6.30 | 26 | -8 | 45 | -10 | Medial Frontal Gyrus | BA 10/11 |
| Right Hemisphere |  |  |  |  |  |  |  |
| 2 | -5.59 | 10 | 10 | -14 | 17 | Thalamus |  |

Note: Temporal lag represents the time after event onset, when a cluster of voxels exhibited a contrast effect of interest. The bootstrap ratio threshold was set to $\pm > 3.5$ , and identified dominant and stable activation clusters.
The spatial extent refers to the total number of voxels included in the cluster (threshold = 10). Gyrus location and Brodmann areas (BAs) were determined by referring to Talairach and Tournoux (1988).

Most of the peaks identified were positive brain saliences. In general, there was a positive correlation between encoding activity in positive salience brain regions and subsequent retrieval accuracy across groups. However in MA_controls_, this effect was only significant during SH encoding events; in MA_+FH_ subjects this effect was only significant during SE encoding events; and in MA_+FH+APOE4_ subjects this was observed for both SE and SH encoding events. At retrieval, activity in these same regions during SE events was positively correlated with memory performance in MA_+FH_, but activity in these regions during SE and SH retrieval was negatively correlated with memory performance in MA_+FH+APOE4_. Therefore, LV1 identified brain regions in which: i) encoding activity was correlated with better subsequent retrieval in MA_controls_; ii) encoding and retrieval activity during SE memory tasks was positively correlated with memory performance in MA_+FH_; and iii) there was phase-related difference between activity and memory performance correlations in MA_+FH+APOE4_. Positive salience brain regions identified in this LV included medial precuneus, bilateral inferior parietal cortex, anterior-medial prefrontal cortex and cingulate, and hippocampus.

##### Brain regions in which there were group differences in brain-behavior correlations at encoding and retrieval between MA_+FH_ vs. MA_+FH+APOE4_ groups

B-PLS LV 2 accounted for 17.94% of the cross-block covariance and identified group differences in brain activity – behavior correlations between MA_+FH_ vs. MA_+FH+APOE4_ groups. This LV identified the effect of having +APOE4 status within the context of having a family history of AD on brain-behavior correlation. Figure 2B presents the singular image and corresponding bar graph depicting the behavior-brain correlation profile for this LV. Table 6 lists the local maxima from this LV. Positive brain saliences listed in Table 6 represent areas in which there was: i) a positive correlation between encoding and retrieval activity and memory performance in the MA_+FH+APOE4_ group, and ii) a negative correlation between encoding activity and subsequent retrieval accuracy in the MA_+FH_ group. These regions primarily included bilateral occipto-temporal cortices and right ventrolateral prefrontal cortex. Negative brain salience regions represent the inverse effect and were regions in which encoding activity was positively correlated with subsequent retrieval in MA_+FH_ subjects, but in which encoding and retrieval activity was negatively correlated with memory performance in MA_+FH+APOE4_ subjects. These regions included primarily anterior medial prefrontal cortex.

**Table 6:**
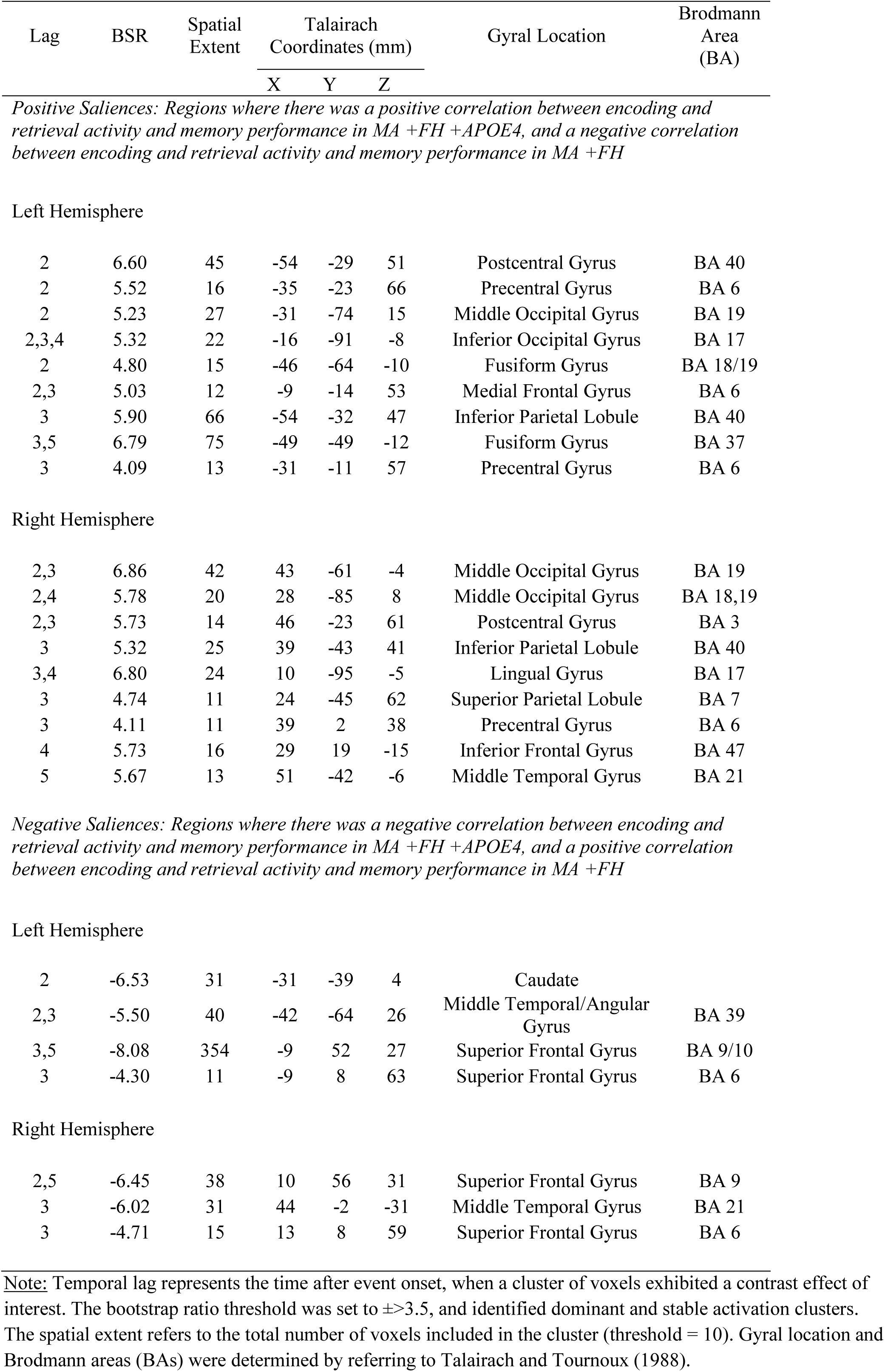
Behavior PLS Results - LV 2

| Lag | BSR | Spatial Extent | Talairach Coordinates (mm) |  |  | Gyrus Location | Brodmann Area (BA) |
| --- | --- | --- | --- | --- | --- | --- | --- |
|  |  |  | X | Y | Z |  |  |
| Left Hemisphere |  |  |  |  |  |  |  |
| 2 | 6.60 | 45 | -54 | -29 | 51 | Postcentral Gyrus | BA 40 |
| 2 | 5.52 | 16 | -35 | -23 | 66 | Precentral Gyrus | BA 6 |
| 2 | 5.23 | 27 | -31 | -74 | 15 | Middle Occipital Gyrus | BA 19 |
| 2,3,4 | 5.32 | 22 | -16 | -91 | -8 | Inferior Occipital Gyrus | BA 17 |
| 2 | 4.80 | 15 | -46 | -64 | -10 | Fusiform Gyrus | BA 18/19 |
| 2,3 | 5.03 | 12 | -9 | -14 | 53 | Medial Frontal Gyrus | BA 6 |
| 3 | 5.90 | 66 | -54 | -32 | 47 | Inferior Parietal Lobule | BA 40 |
| 3,5 | 6.79 | 75 | -49 | -49 | -12 | Fusiform Gyrus | BA 37 |
| 3 | 4.09 | 13 | -31 | -11 | 57 | Precentral Gyrus | BA 6 |
| Right Hemisphere |  |  |  |  |  |  |  |
| 2,3 | 6.86 | 42 | 43 | -61 | -4 | Middle Occipital Gyrus | BA 19 |
| 2,4 | 5.78 | 20 | 28 | -85 | 8 | Middle Occipital Gyrus | BA 18,19 |
| 2,3 | 5.73 | 14 | 46 | -23 | 61 | Postcentral Gyrus | BA 3 |
| 3 | 5.32 | 25 | 39 | -43 | 41 | Inferior Parietal Lobule | BA 40 |
| 3,4 | 6.80 | 24 | 10 | -95 | -5 | Lingual Gyrus | BA 17 |
| 3 | 4.74 | 11 | 24 | -45 | 62 | Superior Parietal Lobule | BA 7 |
| 3 | 4.11 | 11 | 39 | 2 | 38 | Precentral Gyrus | BA 6 |
| 4 | 5.73 | 16 | 29 | 19 | -15 | Inferior Frontal Gyrus | BA 47 |
| 5 | 5.67 | 13 | 51 | -42 | -6 | Middle Temporal Gyrus | BA 21 |
| Left Hemisphere |  |  |  |  |  |  |  |
| 2 | -6.53 | 31 | -31 | -39 | 4 | Caudate |  |
| 2,3 | -5.50 | 40 | -42 | -64 | 26 | Middle Temporal/Angular Gyrus | BA 39 |
| 3,5 | -8.08 | 354 | -9 | 52 | 27 | Superior Frontal Gyrus | BA 9/10 |
| 3 | -4.30 | 11 | -9 | 8 | 63 | Superior Frontal Gyrus | BA 6 |
| Right Hemisphere |  |  |  |  |  |  |  |
| 2,5 | -6.45 | 38 | 10 | 56 | 31 | Superior Frontal Gyrus | BA 9 |
| 3 | -6.02 | 31 | 44 | -2 | -31 | Middle Temporal Gyrus | BA 21 |
| 3 | -4.71 | 15 | 13 | 8 | 59 | Superior Frontal Gyrus | BA 6 |

Note: Temporal lag represents the time after event onset, when a cluster of voxels exhibited a contrast effect of interest. The bootstrap ratio threshold was set to $\pm > 3.5$ , and identified dominant and stable activation clusters. The spatial extent refers to the total number of voxels included in the cluster (threshold = 10). Gyral location and Brodmann areas (BAs) were determined by referring to Talairach and Tournoux (1988).

#### 3 2 3. Post-hoc ROI-based analysis of medial temporal lobe regions identified in PLS analyses

One of our a priori hypotheses was that there would be group differences in MTL activation during spatial context encoding/retrieval. T-PLS LV2 identified two peaks in right hippocampus that were similarly activated during easy > hard encoding across all groups. B-PLS LV1 also identified a region in left hippocampus. To explore if there were between group differences in hippocampal activation we examined group differences in event-related activity within the three hippocampal ROIs identified in T-PLS LV2 and B-PLS LV1 (marked with asterisks in Table 3 and 5). This was done by extracting the mean activity in a 4mm cubic region surrounding each ROI using the multiple voxel extraction option in PLSGUI. We then calculated the mean activity for the ROIs for lags 2 – 5, for each ROI within each subject, and then conducting post-hoc group (3) x event-type (2) x phase (2) repeated measures ANOVAs (significance assessed at p<0.05, corrected).

**Table 3:** Task PLS Results – LV 2

| Lag | BSR | Spatial Extent | Talairach Coordinates (mm) |  |  | Gyrus Location | Brodmann Area (BA) |
| --- | --- | --- | --- | --- | --- | --- | --- |
|  |  |  | X | Y | Z |  |  |
| Positive Saliences: Regions that were more active during hard vs. easy spatial context encoding |  |  |  |  |  |  |  |
| Left Hemisphere |  |  |  |  |  |  |  |
| 3 | 5.08 | 33 | -53 | -60 | 27 | Angular Gyrus | BA 39 |
| 5 | 4.55 | 13 | -12 | -81 | 4 | Lingual Gyrus | BA 17 |
| Right Hemisphere |  |  |  |  |  |  |  |
| 3 | 5.04 | 12 | 54 | -65 | 31 | Angular Gyrus | BA 39 |
| Negative Saliences: Regions that were more active during easy vs. hard spatial context encoding |  |  |  |  |  |  |  |
| Left Hemisphere |  |  |  |  |  |  |  |
| 2,3 | -10.97 | 2071 | -31 | 17 | 5 | Inferior Frontal Gyrus | BA 45 |
| 2,5 | -7.84 | 463 | -9 | 9 | 45 | Medial Frontal Gyrus | BA 32 |
| 2,5 | -5.48 | 68 | -31 | -50 | 39 | Inferior Parietal Gyrus | BA 40 |
| 2,3 | -4.98 | 38 | -34 | 54 | 16 | Superior Frontal Gyrus | BA 10 |
| 2 | -4.77 | 13 | -35 | -85 | 14 | Middle Occipital Gyrus | BA 18 |
| 2 | -4.09 | 11 | -9 | -72 | 30 | Precuneus | BA 31 |
| 2 | -4.80 | 33 | -1 | -34 | 23 | Posterior Cingulate | BA 23 |
| 4,5 | -5.54 | 39 | -49 | -47 | 3 | Middle Temporal Gyrus | BA 21, 37 |
| 4 | -4.12 | 14 | -9 | -17 | 10 | Thalamus |  |
| Right Hemisphere |  |  |  |  |  |  |  |
| 2 | -5.54 | 78 | 13 | -60 | 24 | Precuneus | BA 31 |
| 2 | -4.59 | 63 | 36 | 6 | 34 | Precentral Gyrus | BA 6 |
| 2,5 | -4.15 | 11 | 47 | 34 | 26 | Middle Frontal Gyrus | BA 46 |
| 3 | -5.18 | 30 | 29 | 5 | -24 | Hippocampal Gyrus* | BA 35 |
| 3 | -4.97 | 12 | 40 | -15 | -18 | Hippocampus* |  |
| 4 | -8.94 | 4943 | 6 | 9 | 48 | Superior Frontal Gyrus | BA 6 |

Note: Temporal lag represents the time after event onset, when a cluster of voxels exhibited a contrast effect of interest. The bootstrap ratio threshold was set to $\pm > 3.5$ , and identified dominant and stable activation clusters. The spatial extent refers to the total number of voxels included in the cluster (threshold = 10). Gyrus location and Brodmann areas (BAs) were determined by referring to Talairach and Tournoux (1988). The \* identifies the hippocampal peaks for which post-hoc ROI analyses were conducted.

The post-hoc analysis indicated there was a significant group x task interaction in activity of the right hippocampal ROI (x=40 mm, y =−15 mm, z = −18 mm; F_2,48_=4.73, p<.05det) identified from T-PLS LV2. Figure 2C presents the adjusted mean *%* signal differences in right hippocampus across all events, for each group. To clarify this interaction effect, between-group one-way ANOVAs of ROI activity during eSE, eSH, rSE and rSH, respectively, with post-hoc comparisons on the group variable, were conducted. These one-way ANOVAs indicated there were group differences in encoding activity during eSE events, and no other events. M_controls_ activated right hippocampus to lesser degree than MA_+FH+ApOE4_ subjects (p<0.05). MA_+FH_ subjects’ activation of this region fell midway between the two other groups. In addition, we conducted within phase (encoding, retrieval) x task (SE, SH) group repeated measures ANOVAs of this right hippocampal ROI. These ANOVAs indicated that there was no significant task-related, or phase-related, modulation of right hippocampal activity in either MA_controls_ or MA_+FH_ groups. However, in MA_+FH+APOE4_ there was a significant task main effect in right hippocampal activation (p<0.05) and a trend towards a significant phase x task interaction (p =0.09). Therefore, the group x task interaction was driven by a task-related difference in right hippocampal activation during encoding in MA_+FH+APOE4_.

#### 3 2 4. Post-hoc Hippocampal Volumetric Analysis

To examine whether the aforementioned group differences in hippocampal activity may be related to group differences hippocampal volume, we measured the right and left whole hippocampal volume of each subject using a previously validated automated pipeline called MAGeT (Multiple Automatically Generated Templates brain segmentation algorithm) [43, 44]. Hippocampal volumes were calculated using a multi-atlas voting procedure based on a template library made up of images from the subject pool. The current study used 21 templates and 5 atlases, which were MR images manually traced using a well-established hippocampus segmentation protocol developed by Pruessner [45], which was previously validated in MAGeT [44]. MAGeT output was quality-controlled by an individual who was trained on the same manual segmentation protocol that was used to create the atlases.

The quality control involved scoring each brain for the discrepancy between the MAGeT segmentation and the manual segmentation protocol mentioned above on a slice-to-slice basis [46]. An error was scored each time there was a 20-voxel worth of discrepancy between the MAGeT output and the manual segmentation protocol. Brains that scored sixteen or more on inaccuracy (more than 300 voxels of error) on either the left or right hippocampus, were two standard deviations from the mean, and were manually corrected using the Pruessner [45] protocol to insure the participants’ hippocampi were accurately represented. Since MAGeT yields volumes in native space, we corrected hippocampal volume of each subject by their total brain volume, obtained using the brain extraction based on nonlocal segmentation technique [BEaST],[47]). Manual quality assurance was also performed on these labels.

Two subjects’ automatic segmentation for total brain volume did not pass quality control, and they were excluded from subsequent analysis (1 MA_control_, 1 MA_+FH_). Automatic MAGeT segmentation failed in 2 subjects (1 MA_control_, 1 MA_+_FH) for the left hippocampus and failed in 3 subjects (2 MA_control_, 1 MA_+_FH) in the right hippocampus. These 5 hippocampal volumes were manually corrected by co-author EHY and verified by co-author JCP, and included in the between-group ANOVAs. The total mean volume for left and right hippocampus (HC, mm) unadjusted for total brain volume is presented in Table 1. We calculated the total brain volume (TBV) adjusted HC volume (HC volume/TBV) and conducted one-way ANOVAs to determine if there were significant group differences in HC volume. Between group ANOVAs indicated there was no significant left or right HC volume difference between groups (F<1).

## 4. Discussion

The goal of this study was to use multivariate PLS methods to assess functional brain differences in recollection-related brain activity during the encoding *and* retrieval of spatial contextual details in early middle-aged adults with +FH, and combined +FH, +APOE4, risk factors for late-onset AD, compared to controls. Our behavioral results show there were no significant group differences in spatial context memory ability. In a previous study, we used the same experimental tasks used in the current study to context memory in –FH young adults and –FH MA. We found that there were behavioral deficits on these tasks in MA compared to young adults (Kwon et al., 2016). This suggests that having risk factors for AD did not impact spatial context memory at midlife, beyond the effect of age.

The fMRI results show there were significant group similarities and differences in event-related brain activity and in the correlation between brain activity and retrieval accuracy. The group similarities in event-related brain activity, highlighted in Task PLS LV1 and LV2, indicate that in general, all three MA groups activated similar sets of brain regions during successful encoding and retrieval of spatial contextual details. Successful encoding of spatial contextual details was associated with increased activity in a distributed set of brain regions including: bilateral occipito-temporal cortices, left VLPFC, and bilateral anterior-medial PFC in all three groups, compared to successful contextual retrieval.

These results are consistent with prior studies of face encoding which have also reported increased activity in VLPFC and bilateral occipito-temporal cortices [48–51]. Successful recollection of spatial contextual detail was related to greater activation in bilateral dorsal VLPFC, right DLPFC, left dorsal inferior parietal cortex and bilateral fusiform cortex, compared to encoding. This pattern of retrieval-related activation is consistent with prior studies of episodic retrieval of faces [49, 52, 53].

The group differences in event-related activity and in brain-behavior correlations indicate that even though having +FH and +APOE4 risk factors at midlife did not affect spatial context memory performance; they affected brain activity in memory-related brain regions, and altered patterns of brain-behavior associations. More specifically, there were three main group differences in event-related activity and brain-behavior correlations identified in the current study. First, both MA groups with AD risk factors exhibited greater activation in hippocampus during SE encoding events compared to MA_controls_, and increased hippocampal activity at encoding correlated with better subsequent memory performance. Second, there were group differences in activity and brain-behavior correlations in left angular gyrus, cingulate gyrus and precuneus in MA with AD risk factors, compared to controls. Third, activity and brain activity-behavior correlations in anterior-medial PFC and in ventral visual cortex differentiated the two MA risk groups from each other, and from MA_controls_. We discuss these group differences in brain activity and brain-behavior correlation in detail in the following sections.

### 4. 1 Group differences in hippocampal activity

T-PLS LV2 results indicated there was increased right hippocampal activation during easy > hard spatial context encoding across all groups. This is surprising since prior task fMRI studies have reported altered MTL activity in adults at genetic risk of developing AD [27, 54, 55] and have indicated there is an interaction between +FH and +APOE4 risk factors on MTL activity [20, 21]. However, these studies used univariate and region-of-interest analysis methods; not the data driven multivariate approach used in the current study. Indeed, when we conducted post-hoc univariate analyses of the hippocampal peaks identified by PLS, we observed group differences in right hippocampal activation. Specifically, MA_+FH+APOE4_ subjects exhibited greater activity in right hippocampus during SE encoding events compared to other event-types, and the level of activity they exhibited was significantly greater than that observed in MA_controls_. MA subjects with only a +FH risk factor exhibited activation levels in right hippocampus during SE encoding that were midway between MA_controls_ and MA_+FH+ApoE4_ subjects. Therefore, our univariate analysis indicated that MA with risk factors for AD over-activated the hippocampus during SE encoding, compared to MA controls. Interestingly, this difference in activation was apparent even though there were no group differences in hippocampal volume, nor group differences in performance.

We also observed a positive correlation between encoding activity in left hippocampus and subsequent retrieval in all three groups (B-PLS LV1). This indicates that the increased hippocampal activity observed in MA with AD risk factors, compared to controls, supported memory. This interpretation is consistent with the observation that MA with AD risk factors over-activated this region during SE encoding, compared to controls – the encoding task which was related to better subsequent retrieval, and thus reflected more successful encoding, compared to the SH task. Therefore, our results show that: 1) MA with AD risk factors exhibited greater hippocampal activity, compared to controls, during SE encoding; 2) hippocampal activity during SE encoding was correlated with better subsequent memory; and, 3) there were no group differences in memory performance in the current study. Taken together these results are consistent with the hypothesis that over-recruitment of hippocampal activity in MA at risk of AD during SE encoding tasks reflects “successful” compensation at encoding [56–58]. However, it is important to note that our post-hoc volumetric analysis of the hippocampus did not identify significant group differences. Therefore, over-activation of the hippocampus was apparent in the absence of underlying gray matter volume loss in MA with AD risk factors, compared to controls. Yet, it is important to note, that this negative volumetric findings may be due to the small sample size in the current study and a lack of statistical power to detect group differences in hippocampal volume.

The current findings are in contrast to Trivedi et al (2006) who reported MA with +FH and +APOE4 AD risk factors exhibited *less* activity in hippocampus compared to MA with only the +FH risk factor or neither risk factors (controls). Trivedi et al (2006) also reported a positive association between left medial temporal activity during encoding and performance on the Rey Auditory Learning Task. However, there were several differences between the methods employed by Trivedi et al (2006) and the current study. First, Trivedi et al (2006) tested adults who were on average older than our sample [19]. Given that hippocampal activity has been shown to decrease with age during episodic encoding; it is possible that the difference between our current results and Trivedi et al. is due to the differing age of the groups tested. This interpretation is consistent with the observation that our findings are more similar to those reported by Filippini et al (2009) and Dennis et al (2010) who reported increased hippocampal activation at encoding in *young* +APOE4 carriers [54, 55]. In fact, Filippini et al (2011) showed that there may be interaction between age and APOE4 impact on hippocampal activity [28]. Thus, testing on average younger (40-58yrs, in this study) vs. older (40-65, Trivedi et al (2006)) middle-aged adults may have affected the pattern of hippocampal activation observed in our study compared to Trivedi et al (2006).

Second, Trivedi et al (2006) employed an item memory task to examine brain activation during novel (item encoding), compared to repeated (familiar), presentations of object stimuli across groups. All subjects in this task scored at ceiling (98% or higher) for novelty detection. Therefore, it is unclear if novelty detection in the study by Trivedi et al (2006) reflected encoding success, as measured in the current study, since Trivedi et al (2006) did not conduct a post-scanning retrieval test for objects presented in the scanner. Moreover, in the current study subjects were aware their memory for spatial context would be tested subsequently. Thus, the experimental design and task demands were significantly different between studies. This highlights a key issue about the interpretation of group differences in hippocampal activity during incidental encoding, as measured with novelty detection, compared to intentional encoding, as measured in the current study. One possibility is that MA with AD risk factors do not automatically activate the hippocampus to support incidental encoding, whereas MA controls do (Trivedi et al, 2006). However, when directed to perform a memory task, MA with AD risk factors are able to intentionally activate the hippocampus, and do so to a greater degree than MA controls during low demand conditions (current study). This pattern of association between hippocampal activity and behavior is reminiscent of studies of frontal lobe function in cognitive aging [59–62]. This literature has shown that older adults do not spontaneously engage frontal-related strategic processes to assist memory encoding to the same degree as young adults. However, during intentional encoding conditions, older adults over-activate the frontal lobes at encoding compared to young adults – especially when performance was matched between groups [59, 63]. These findings have been taken to support they hypothesis that there may be subtle deficits in frontal function with age that impede episodic memory, and that older adults over-activate the frontal lobes during intentional encoding compared to young adults to compensate for these subtle deficits. We propose a similar interpretation can be applied to explain the discrepant findings by Trivedi et al (2006) and the current study: that MA with AD risk factors are showing subtle deficits in hippocampal function, which they successfully compensate for by over-activating this region during intentional encoding tasks. This conclusion is supported by results from Bassett et al (2006). In this study, adults aged 50-75 yrs, with +FH and +APOE4 risk factors, exhibited greater activity in hippocampus compared to controls during an intentional encoding study for verbal paired associations. Therefore, task demands may affect whether MA with AD risk factors exhibit decreased or increased MTL activity compared to controls.

### 4.2. Group differences in brain activity and/or in brain activity-behavior correlations in cortex

T-PLS LV3 identified group differences in task-related activity in a variety of brain regions including: inferior parietal cortex, cingulate gyrus, precuneus and ventral fusiform cortices. Importantly, these activations overlapped with areas identified in the B-PLS results. In addition, there were activations identified in the B-PLS results (i.e. medial PFC), which overlapped with activations in T-PLS LV1, which highlight group similarities in encoding vs. retrieval activity. In the following subsections we discuss brain regions than were identified in both T-PLS and B-PLS results to better understand how AD risk factors impact both activity and brain-behavior associations at midlife.

#### 4.2.1. Overlap in regions identified in T-PLS LV3 and B-PLS LV1

Bilateral inferior parietal cortex, cingulate gyrus and precuneus were positive salience areas from T-PLS LV3. This indicates these regions were more active during encoding, compared to retrieval, in MA_controls_ and MA_+FH_ subjects, and were more active during the SE retrieval, compared to SE encoding, in MA_+FH+APOE4_ subjects. These brain regions were also positive salience areas in B-PLS LV1. Thus, in MA_controls_, encoding activity in these brain regions was positively correlated with subsequent recollection, particularly during SH events (see Figure 2a). In MA_+FH_ subjects, encoding activity in these regions was positively correlated with subsequent recollection of SE events; but the pattern of retrieval activity was negatively correlated with recollection of SE events. In MA_+FH+APOE4_ subjects the pattern of event-related activity observed in bilateral inferior parietal, cingulate gyrus and precuneus was not beneficial to their task performance. Specifically, the B-PLS LV1 correlation profile (Figure 2a) indicates that increased encoding activity and decreased retrieval activity in these areas was positively correlated with memory performance for this group. However, the Task PLS result (Figure 1b) shows that this group exhibited the opposite pattern of activation in these brain regions: increased activity during retrieval, and decreased activity during encoding of SE events. Taken together these results indicate that there may be a progressive difference in brain activity and brain-behavior correlations involving the inferior parietal cortex, cingulate gyrus, and precuneus going in MA_+FH_ and MA_+FH+APOE4_, compared to controls.

**Figure 1:**
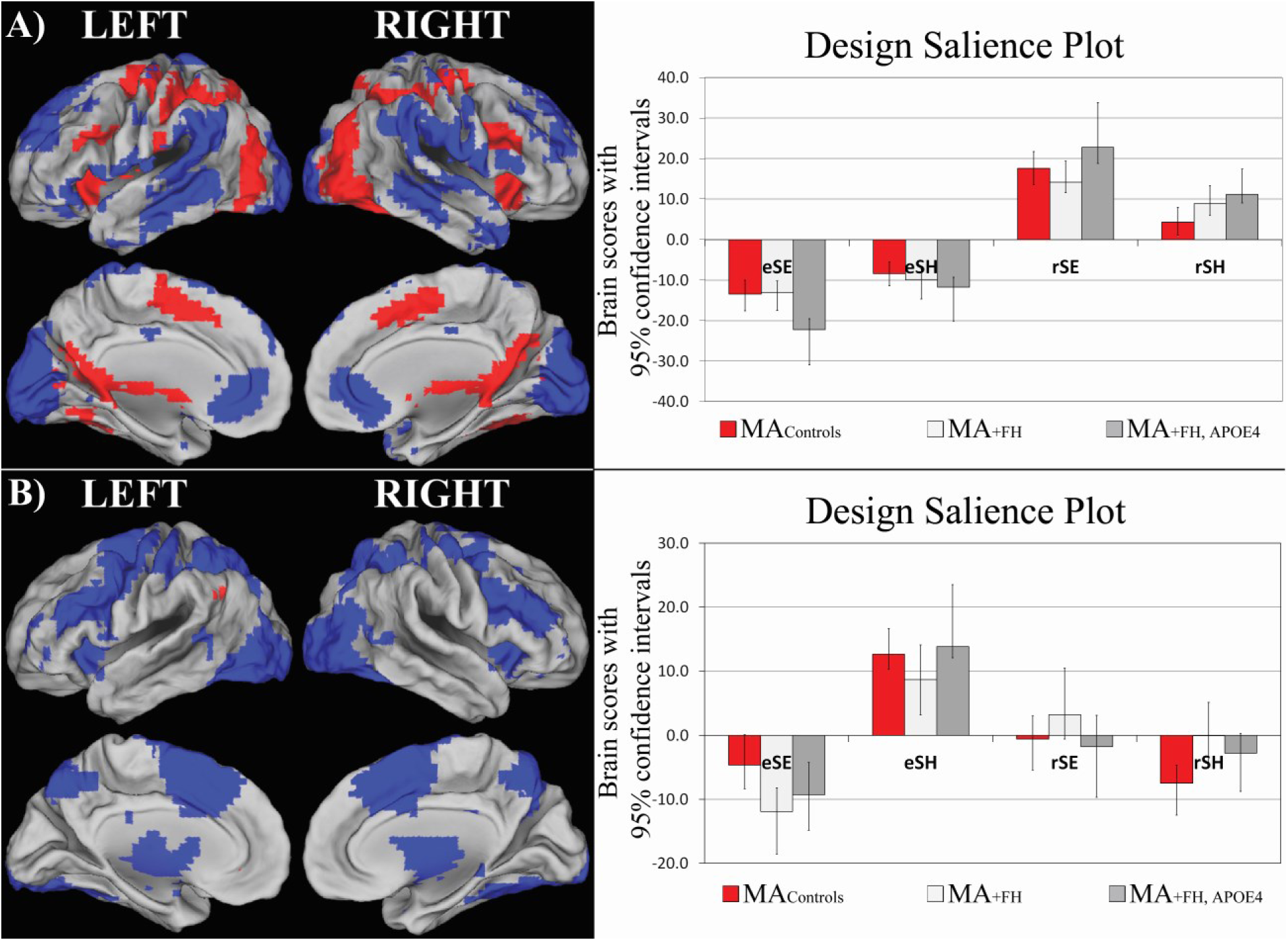
T-PLS LV1 and LV2 Result. A) The singular image and design salience plot for T-PLS LV1. The singular image is thresholded at a bootstrap ratio of ±3.5, p<0.001. Red brain regions reflect positive brain saliences and blue regions reflect negative brain saliences. Activations are presented on template images of the lateral and medial surfaces of the left and right hemispheres of the brain using Caret software (http://brainvis.wustl.edu/wiki/index.php/Caret:Download). The design salience plots represent the brain scores with 95% confidence intervals (y-axis) for each group for each task-type (x-axis). eSE = encoding, easy spatial context memory tasks; eSH = encoding, hard spatial context memory tasks; rSE = retrieval, easy spatial context memory tasks; rSH = retrieval, hard spatial context memory tasks. The design salience plot for T-PLS LV1 indicates this LV identified brain regions that were differentially activated during successful spatial context retrieval (positive saliences) vs. encoding (negative saliences). B) The singular image and design salience plot for T-PLS LV2. The singular image is thresholded at a bootstrap ratio of ±3.5, p<0.001. This LV identified brain regions that were differentially activated during hard spatial encoding (eSH; positive saliences) vs. easy spatial encoding (eSE; negative saliences).

**Figure 2:**
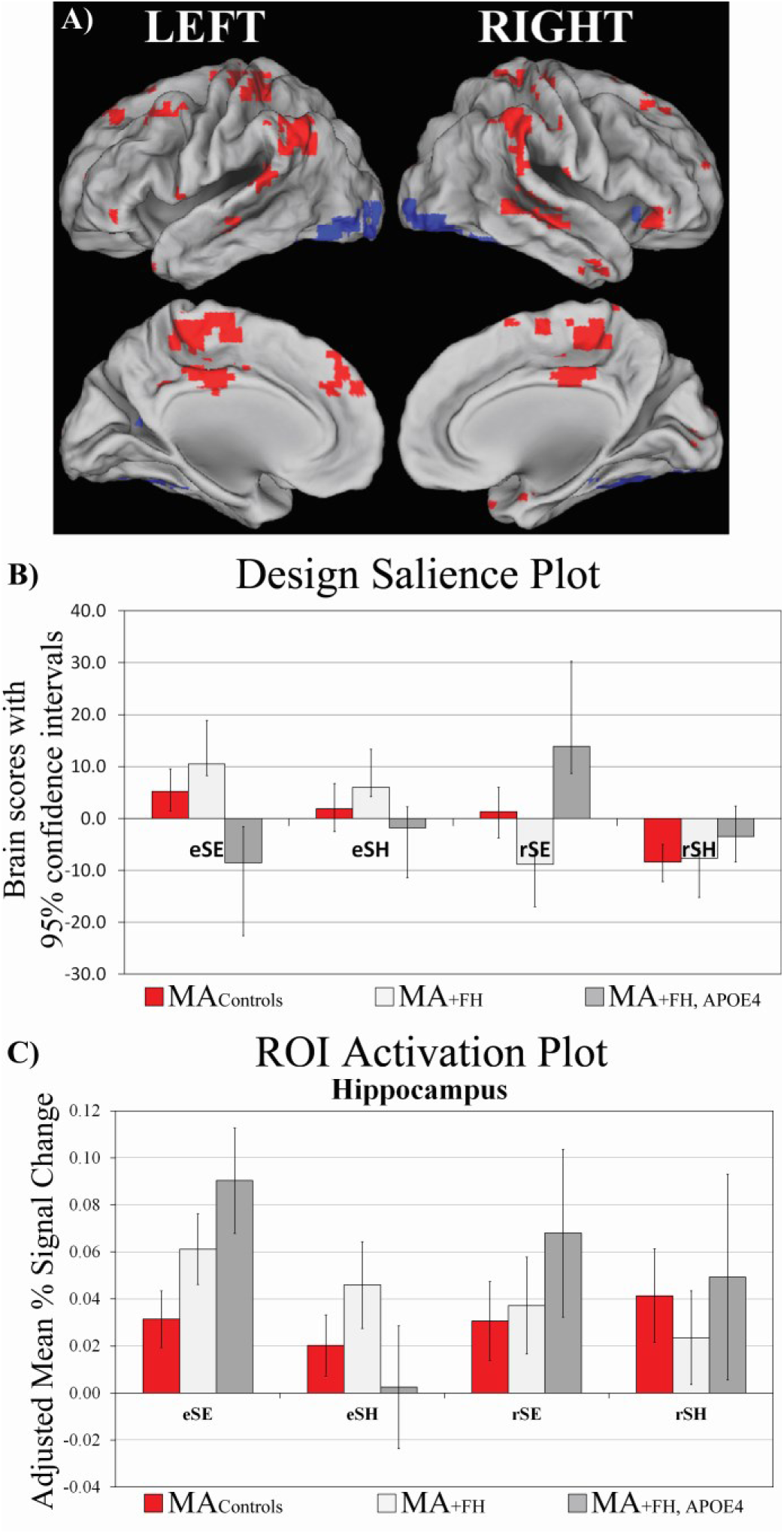
T-PLS LV3 and Right Hippocampal Activation Plot. **A**) The singular image and **B**) design salience plot for T-PLS LV3. The singular image is thresholded at a bootstrap ratio of ±3.5, p<0.001. Red brain regions reflect positive brain saliences and blue regions reflect negative brain saliences. The design salience plot for T-PLS LV3 indicates this LV identified a group^*^task^*^phase interaction. Positive salience brain regions were more active during encoding > retrieval in MA_+FH_ and MA_controls_; and more activity during easy spatial context retrieval > easy spatial context encoding in MA_+FH+APOE4_. Negative salience brain regions exhibited the inverse pattern of associations. C) ROI Activation Plot for right hippocampus from T-PLS LV2.

**Figure 3:**
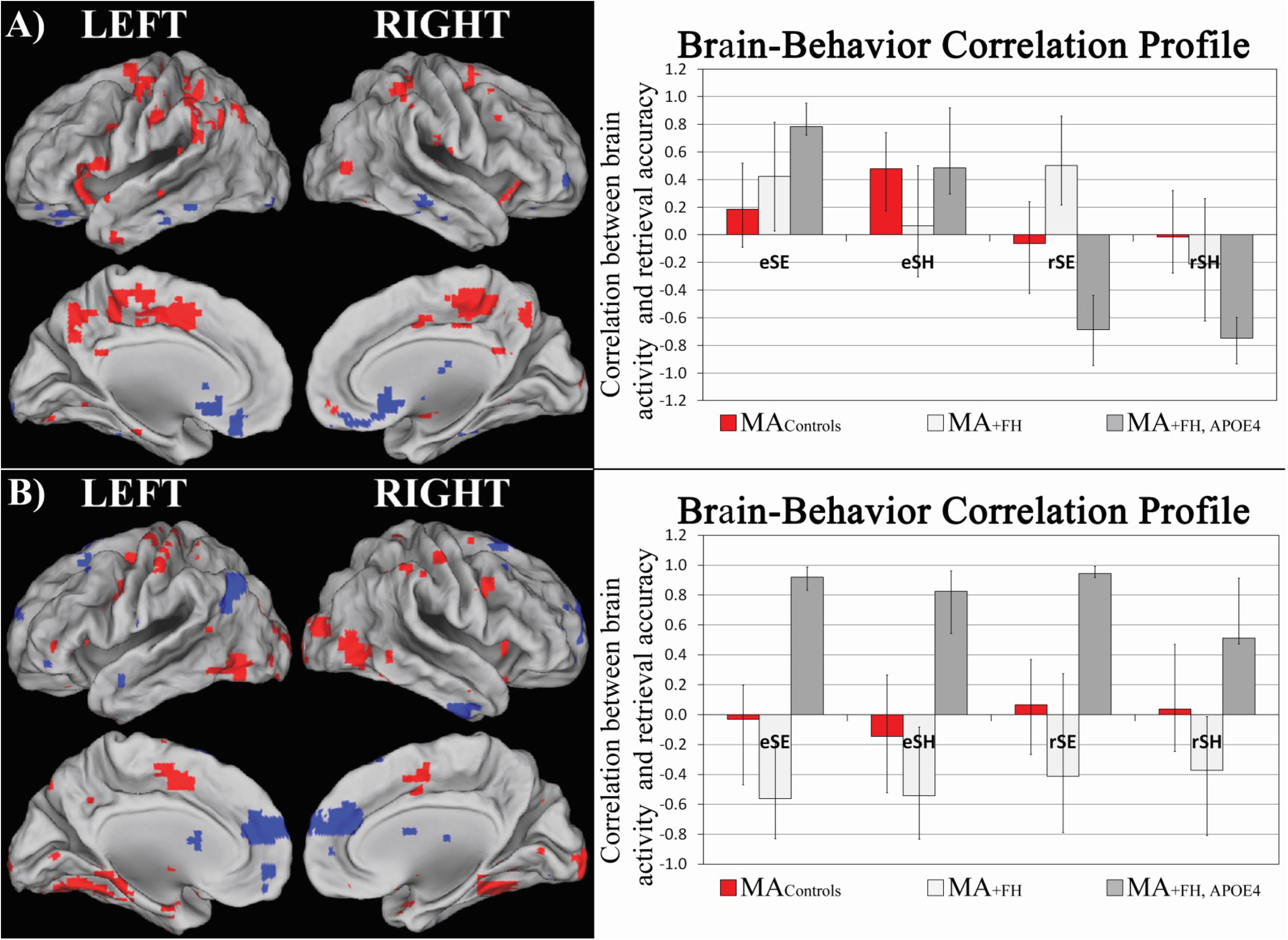
B-PLS Results. A) The singular image at a bootstrap ratio threshold = ±3.5, p<0.001 and the brain-behavior correlation profile with 95% confidence intervals for B-PLS LV1. In the singular image red brain regions reflect positive brain saliences and blue regions reflect negative brain saliences. Activations are presented on template images of the lateral and medial surfaces of the left and right hemispheres of the brain using Caret software. The correlation profile indicates that encoding activity in positive salience regions was positively correlated with subsequent retrieval for both task in MA+FH+APOE4 and MAcontrols; and retrieval activity in these same regions was negative correlated with retrieval accuracy on both tasks in MA_+FH+APOE4_. In MA+FH encoding and retrieval activity in positive salience regions during SE tasks was correlated with better performance on this task. Negative salience regions exhibited the inverse pattern of brain-behavior correlations. **B)** The singular image at a bootstrap ratio threshold = ±3.5, p<0.001 and the brain-behavior correlation profile with 95% confidence intervals for B-PLS LV2. The correlation profile indicates that increased encoding and retrieval activity in positive salience regions was positively correlated with memory performance on both tasks in MA_+FH+APOE4_ subjects, and negatively correlated with memory performance on both tasks in MA_+FH_. Negative salience regions exhibited the inverse pattern of brain-behavior correlations.

Interestingly, inferior parietal, cingulate gyrus and precuneus are key nodes of the default mode network (DMN), which is a functionally connected set of brain regions found to be more active during baseline vs. task conditions in fMRI studies [64–66]. Several studies have reported differences in DMN activity and functional connectivity in MCI, AD, and in healthy adults with risk factors for AD [67–71]. The fact that we observed memory-related activations in inferior parietal, cingulate and precuneus, is not surprising since previous studies have noted the overlap in brain activation patterns during autobiographical/episodic memory processing and resting state [72–75]. It has been hypothesized that these regions may be involved in the attentional processing and integration of one’s experience, which occurs during both rest and episodic memory task performance [75–78]. Although, in the current study we did not observe overt behavioral deficits in MA with AD risk factors, compared to controls; the current B-PLS results suggest that there may be subtle, differences in inferior parietal, cingulate and precuneus function in both MA groups with AD risk factors, compared to controls, which was negatively related to their memory performance. This suggests that having a family history of AD may alter the function of these brain regions at midlife, since this risk factor was common to both MA risk groups.

#### 4.2.2. Overlap in regions identified in T-PLS LV3 and B-PLS LV2

T-PLS LV3 identified significant group differences in bilateral fusiform cortex activation. More specifically, these regions were more active during retrieval, compared to encoding in MA_controls_ and MA_+FH_ subjects. In contrast, these regions were more active during SE encoding, compared to SE retrieval, in MA_+FH+APOE4_ subjects (see Figure 1). These brain regions were also positive salience areas in B-PLS LV2 (see Figure 2B). Therefore, MA_+FH_ adults had decreased activity in fusiform cortex at encoding, compared to retrieval; and, this was positively related to subsequent retrieval. MA_+FH+APOE4_ adults had increased activity in this same region at encoding compared to retrieval, and this was positively correlated with subsequent retrieval. In contrast, at retrieval, MA_+FH+APOE4_ exhibited decreased fusiform activity, relative to encoding; and this pattern of retrieval activity was negatively correlated to memory performance. MA_+FH_ exhibited increased activity in ventral visual regions at retrieval, compared to encoding, but this was not strongly associated with memory performance.

Overall, these results show that MA_+FH+APOE4_ differentially activated fusiform cortex during encoding and retrieval, compared to MA_+FH_. However, by combining T-PLS with B-PLS results we see that even though the two MA risk groups displayed distinct activation profiles for these regions, the impact on memory performance was similar across both groups. One speculative interpretation is that these differences in fusiform activity at encoding may reflect the utilization of distinct encoding strategies in each of the two at-risk groups, respectively, to support spatial context encoding. For example, in our prior work on healthy aging using the same paradigm we have reported that successful spatial context memory in young adults was associated with increased activity in fusiform cortex[12, 79]. We interpreted this as reflecting young adults’ vivid encoding of perceptual details, which supported subsequent memory. Others have also reported that increased ventral visual activity at encoding and retrieval in young adults supported vivid encoding and detailed recollection [80, 81].

This suggests that MA_+FH+APOE4_ adults may be relying on similar perceptual strategies to support spatial context encoding in the current study. In contrast, MA_+FH_ may be using a non-perceptual strategy, i.e. socio-affective strategy at encoding – similar to what we have reported in older adults in this same paradigm [12, 79] (discussed below). Thus, decreased activity in fusiform cortex at encoding may have supported their utilization of a non-perceptual strategy. Additional research is needed to determine if altered ventral visual function is consistently observed in adults with both +FH and +APOE4 risk factors and how it relates to memory performance.

#### 4.2.3. Group differences in brain-behavior correlations involving anterior-medial PFC

Anterior-medial PFC was identified as a negative brain salience region in T-PLS LV1 and in B-PLS LV2. T-PLS LV1 identified group similarities in phase-related differences in brain activity at encoding and at retrieval, and indicated that all groups exhibited greater activity in anterior-medial PFC at encoding compared to retrieval. B-PLS LV2 indicated that increased encoding activity in anterior-medial PFC was correlated with better subsequent memory performance in MA_+FH_ adults, and poorer subsequent memory performance in MA_+FH+APOE4_ adults. Encoding activity in this region was not significantly correlated with performance in MA_controls_.

Prior fMRI studies of adults with AD risk factors have reported disruptions in the activation and functional connectivity of the anterior-medial PFC [67, 70, 71]. Moreover, generalized increased activity in medial PFC has been observed in AD patients and older adults with mild cognitive impairment compared to controls [82, 83]. The current results indicate that having AD risk factors did not impact the pattern of anterior-medial PFC activity at midlife, but there was an interaction in how +FH and +APOE4 risk factors impacted how anterior-medial PFC activity correlated with memory performance.

In the current study, at encoding, subjects had to make pleasant/neutral judgements for face stimuli while simultaneously encoding the face-location association. Activity in anterior-medial PFC at encoding has been associated with the use of subjective encoding strategies [84, 85]. Thus, our T-PLS results suggest that greater anterior-medial PFC activity at encoding may reflect subjects’ engaging subjective value-based processes to make the pleasantness judgments of face stimuli at encoding. Interestingly, our B-PLS results suggest that MA_+FH_ subjects’ memory performance benefitted from processing the subjective (pleasantness) aspects of the face stimuli during spatial context encoding. This may reflect a shift in using subjective vs. objective stimulus information to make a retrieval judgment in MA_+FH_⋅ Similar results have been reported in fMRI studies of episodic memory in healthy older, compared to younger, adults [86–88]. Therefore, MA_+FH_ may be exhibiting this shift in processing preferences at an earlier age, compared to other MA groups; or this may be a group difference in memory processing in general, which is unrelated to aging.

In contrast, MA_+FH+APOE4_ subjects exhibited a negative correlation pattern between anterior-medial PFC activity and memory performance. This correlation pattern is similar to what has been reported in a meta-analysis of subsequent memory effects in healthy young adults [89]. In young adults, activation of anterior-medial PFC during encoding has been associated with poorer subsequent memory when the memory tasks employed required subjects to encode objective stimulus information[89]. This suggests that for MA with both AD risk factors, successful memory performance was related to successful encoding of objective vs. subjective information, similar to what is observed in young adults [63, 86, 87]. This is consistent with the positive correlation observed between ventral visual encoding activity and memory performance in both MA groups with AD risk factors (discussed above).

### 4.3 Caveats

Despite the strengths of the current study, there are some caveats to our findings. First, we have relatively small sample sizes for the MA_+FH+APOE4_ risk groups which may affect the generalizability of the current findings. However, the sample size used in the current study is comparable to previously published work [21, 25, 54, 90, 91]. Also, we used multivariate PLS with permutation and bootstrap methods to assess the statistical significance and stability of our fMRI results, which are robust methods that are more amenable for use with smaller sample sizes compared to more traditional univariate fMRI methods [92, 93]. Another caveat is that we did not have a –FH, +APOE4 MA sample, since only a small portion of the general population are +APOE4 [94]. This prevented us from differentiating the unique impacts of +FH vs. +APOE4 risk factors on the neural correlates of spatial context memory further.

## 5. Conclusions

The current study examined group similarities and differences in brain activity and brain-behavior correlations during spatial context memory encoding retrieval in MA with vs. without +FH and +APOE4 risk factors for late-onset sporadic AD. Overall, we found no significant group differences in: spatial context memory retrieval accuracy, hippocampal volume, and the general patterns of brain activity associated with successful spatial context encoding and retrieval. However, we also observed group differences in activity and brain-behavior correlations in left inferior parietal cortex, cingulate and precuneus; and group differences in hippocampal activity during encoding, in MA with AD risk factors, compared to controls. This suggests that having +FH or +FH and +APOE4 risk factors alters the activity and brain-behavior correlations in brain regions associated with successful episodic memory encoding and retrieval of spatial contextual details by midlife. In addition, we found that activity and brain activity-behavior correlations in anterior-medial PFC and in ventral visual cortex differentiated the two MA risk groups from each other, and from MA_controls_. This suggests that there is an interaction between +FH and +APOE4 risk factors, such that having both risk factors differentially impacts the memory-related functions in anterior-medial PFC and ventral visual cortex. Overall, the current study broadens our understanding of how having +FH only vs combined +FH, +APOE4 AD risk factors alter brain activity and brain-behavior associations at midlife, and identifies subtle changes in the functional neuroanatomy of episodic memory in middle-aged adults with APOE4 and FH risk factors for AD.

## acknowledgements

This work was supported by the Canadian Institutes of Health Research (CIHR) – operating grant (Grant No. 126105) and Alzheimer’s Society of Canada Research Program Grant (Grant No. 1435) awarded to MNR. LMKW was supported by a STOP-AD fellowship and McGill Graduate Student Excellence Award. We thank Dr. J. Breitner and members of the STOP-AD Centre, McGill University, for subject referrals. The authors declare no conflicts of interest.

